# Neural Trajectories of Conceptually Related Events

**DOI:** 10.1101/2023.12.04.569670

**Authors:** Matthew Schafer, Philip Kamilar-Britt, Vyoma Sahani, Keren Bachi, Daniela Schiller

## Abstract

In a series of conceptually related episodes, meaning arises from the link between these events rather than from each event individually. How does the brain keep track of conceptually related sequences of events (i.e., conceptual trajectories)? In a particular kind of conceptual trajectory—a social relationship—meaning arises from a specific sequence of interactions. To test whether such abstract sequences are neurally tracked, we had participants complete a naturalistic narrative-based social interaction game, during functional magnetic resonance imaging. We modeled the simulated relationships as trajectories through an abstract affiliation and power space. Using two independent samples (n = 50), we found evidence of both the underlying social dimensions of affiliation and power and the evolving social relationships themselves as being tracked by the hippocampus. These results suggest that our evolving relationships with others, despite being composed of isolated events that occur independently across time and space (i.e., spatially and temporally non-consecutive events), but are nevertheless related conceptually, are represented in trajectory-like neural patterns in the human hippocampus.

## Introduction

When you are in a certain place, say a hotel lobby, the meaning of that place (i.e., your location in state space, or latent state) only makes sense in the context of what led you there. A sequence of observations is necessary to disambiguate the current state, as a single observation is insufficient: checking in and checking out of a hotel may look identical yet they predict very different futures. This also applies to conceptual spaces, where sensory information is fully abstracted out. A prototypical case of conceptual space is social space, where the state of a social relationship evolves over interactions along the latent dimensions of power and affiliation^1^. For example, a conversation between two people in that hotel lobby can mean different things if they are old friends, lifelong enemies, or strangers. The prior relationship state—itself a summary of conceptually connected events from the relationship history—helps determine both the current and future relationship states. But while neural representations of latent states and paths in physical space have been identified^2,3^, as have location-like representations in conceptual spaces^4,5^, including social^1,6^, it is unknown whether the brain dynamically tracks locations as a sequence of states in conceptual space.

Where and how could relationship trajectories through social space be represented in the human brain? The hippocampus is a likely candidate. Functional magnetic resonance imaging (fMRI) studies have shown that the hippocampus tracks latent states across domains, from physical locations,^7^ to concepts,^8,9^ to abstract relations between self and other^6,10,11^. Sequences of neural states in the hippocampus may also be connected like trajectories. Evidence consistent with this idea showed that post-task fMRI activity patterns reactivated sequentially^12^, reflecting previously learned sequences. An intriguing possibility is that the hippocampus maps sequences of social interactions in relationships onto a neural manifold, tracking them like trajectories in an abstract social space.

To examine this, we used representational similarity analyses^13^ to test whether hippocampal fMRI patterns show structures that are specific to different relationships’ trajectories, but that share the same representational geometry. Participants completed a naturalistic and narrative-based social interaction task^6^, where they interact and form relationships with fictional people in a new social network. The interactions are defined by choices with latent affiliation and power content, and the relationships with the characters evolve on these dimensions as the task progresses (see **Figure 1**). We tested the following hypotheses about hippocampal representations: First, we expected the hippocampus to represent the social interactions on affiliation (e.g., cooperation) and power (e.g., hierarchy) dimensions, a compressed format that allows generalization across social situations. Across the different characters and interactions, hippocampal patterns should be more similar for interactions of the same dimension than interactions of a different dimension. But while these dimensions are general, the individual relationships are specific: each relationship should have hippocampal representations that reflect the trial-by-trial behavioral trajectories in a shared social space. Our results support these hypotheses and suggest that the hippocampus tracks sequences of relationship-specific social interactions like trajectories on a common manifold. These results provide evidence for hippocampal linking of conceptually related events, even when separated in time and context, into coherent narratives.

**Figure 1.**
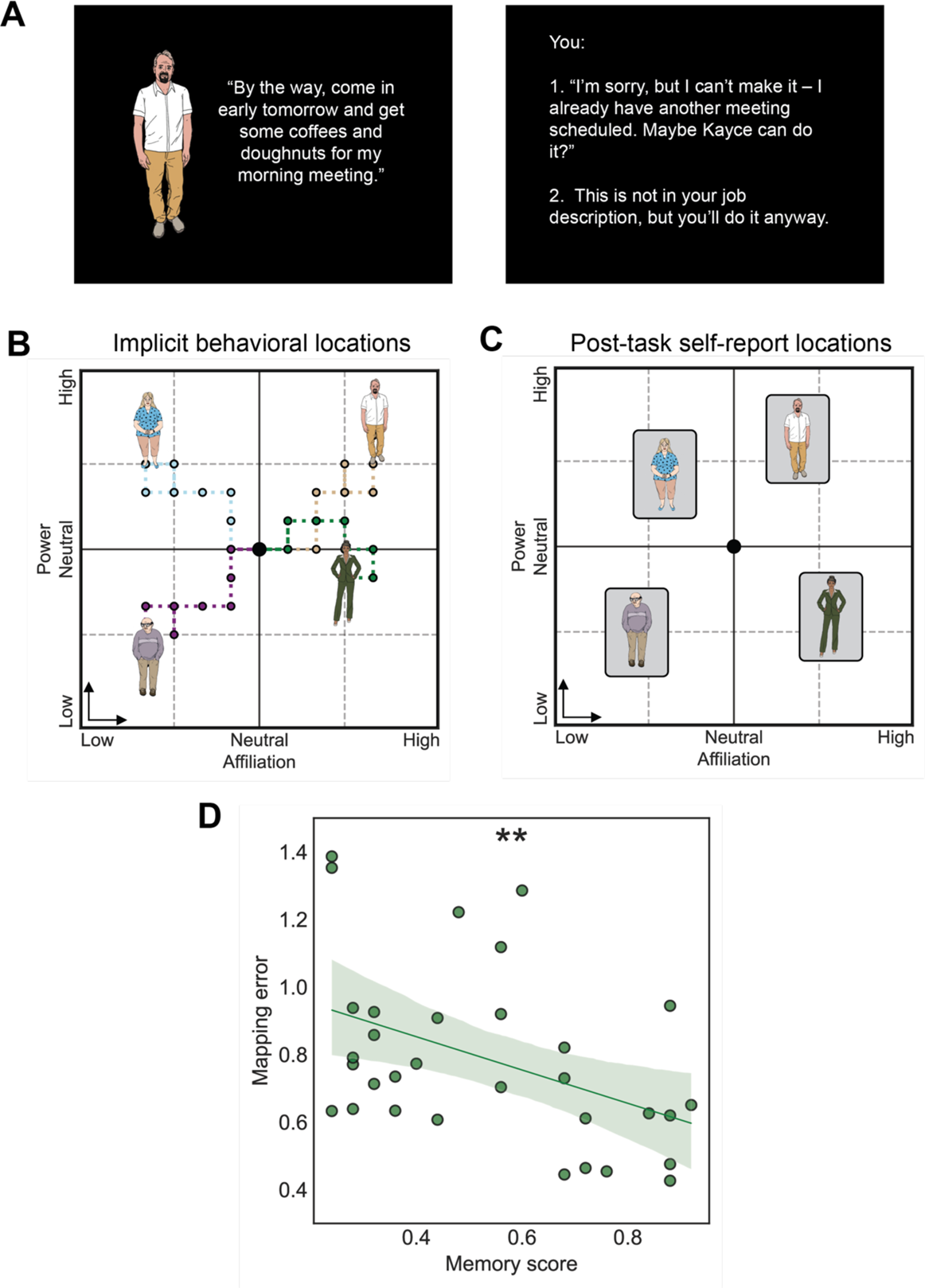
Social interaction sequences form relationship trajectories along abstract dimensions of affiliation and power. (A) An example of a power interaction. Participants read text that describes the narrative and on decision trials choose between two options. Based on their choice, the character moves −1 or +1 along the active dimension (power or, on other trials, affiliation). (B) The participant forms relationships with different characters through sequences of interactions in the narrative. The decisions the participant makes in the implicit affiliation and power interactions change the character’s location in social space, forming a relationship trajectory (participants are unaware of the dimensions). Participants interact with 6 characters: 5 each with 6 affiliation and 6 power trials and 1 with 3 neutral trials. 4 characters are shown for illustration. (C) Schematic of post-task self-report placements. The mapping error between the behavioral and subjective placements was calculated as the average character-wise Euclidean distance between the locations; this was significantly smaller than expected by chance. (D) Mapping error was negatively correlated with task memory, suggesting the subjective maps depend on memory. 95% confidence intervals for regression line are indicated by the shaded region and p-value significance is indicated by asterisks: * < 0.05. Validation sample only (n = 32).

## Results

### Behavioral geometry is consistent with social mapping

Before running the fMRI analyses, we asked whether participants’ choices were consistent with a map-like representation in affiliation and power space. We tested two main behavioral assumptions: (1) behavior is independent between participants (i.e., not a task artifact shared across participants) and (2) behavioral locations align with participants’ self-reported placements of the characters (**Figure 1**). Below, we discuss each of these.

### Participants’ behavior is idiosyncratic

If the task structure determined behavior, participants would have made similar choices and traversed similar paths through social space. However, each participant’s set of choices was unique (i.e., never identical between participants). The character specific trajectories were also largely unique: out of the character specific choices (250 for 50 participants with 5 characters each), 96.8% (242/250) were unique. Moreover, the behavioral trajectories approximately occupied the entire space across both samples. As such, participant behavior was idiosyncratic and unlikely to simply reflect task structure.

### Behavioral locations are related to self-reported locations

To test if the participants subjectively represented the social locations implied by their behavioral choices, we asked whether self-reported (i.e., explicit) placements of the characters into the social space align with the behavioral (i.e., implicit) locations from the task. After the task we had the Validation sample participants (n = 32) place the characters into a two-dimensional (2D) affiliation and power space based on their perception of their relationships with the characters. We calculated the participant-specific average distance between the characters’ behavioral and the self-reported locations (i.e., “mapping error”) and compared them to participant-specific permutation generated distances to get z-scores. A one-sided t-test with these z-scores showed that the distances were smaller than participant-specific permutation-based chance (*β* = −0.563, CI_95_ = [−0.935, −0.191], *t*_31_ = −3.09, *p* < 0.005). We also found a predicted negative correlation between mapping error and task memory (*r* = −0.44, CI_95_ = [−0.767, −0.108], *t*_31_ = −2.71, left-tailed *p* < 0.01), consistent with participants using their memory of the relationships to place the characters in social space. Together these results suggest that affiliation and power locations reflect a subjectively accessible map of the relationships, supporting the internal validity of the task.

### The hippocampus represents the dimensions of affiliation and power during social interactions

After validating our main behavioral assumptions, we turned to the fMRI data. We hypothesized that the social interactions are represented abstractly along the affiliation and power dimensions in the hippocampus, such that they generalize across characters and interactions. If true, decision trial pairs of the same dimension should have more correlated patterns than decision trial pairs of different dimensions: the average pattern correlations should be higher for same dimension versus different dimension decision trial pairs. We used a representational similarity analysis (RSA) searchlight to test this prediction. For each participant, we computed the correlation differences between the same and different dimension trial pairs using a moving searchlight. Then, in the group-level model, we controlled for sample and the average reaction time difference between affiliation and power decisions and tested for a positive effect of correlation difference.

We first tested this in the bilateral hippocampus. There was a significant cluster, with the peak voxel in the anterior hippocampus (peak voxel MNI x/y/z = −34/−17/−16, cluster extent = 724 voxels), supporting our a priori hypothesis that the hippocampus represents social situations along the dimensions of affiliation and power (see **Figure 2**). Importantly, character identity and temporal distance (i.e., the time between trials) are experimentally balanced across affiliation and power decision trials, and cannot explain these effects.

**Figure 2.**
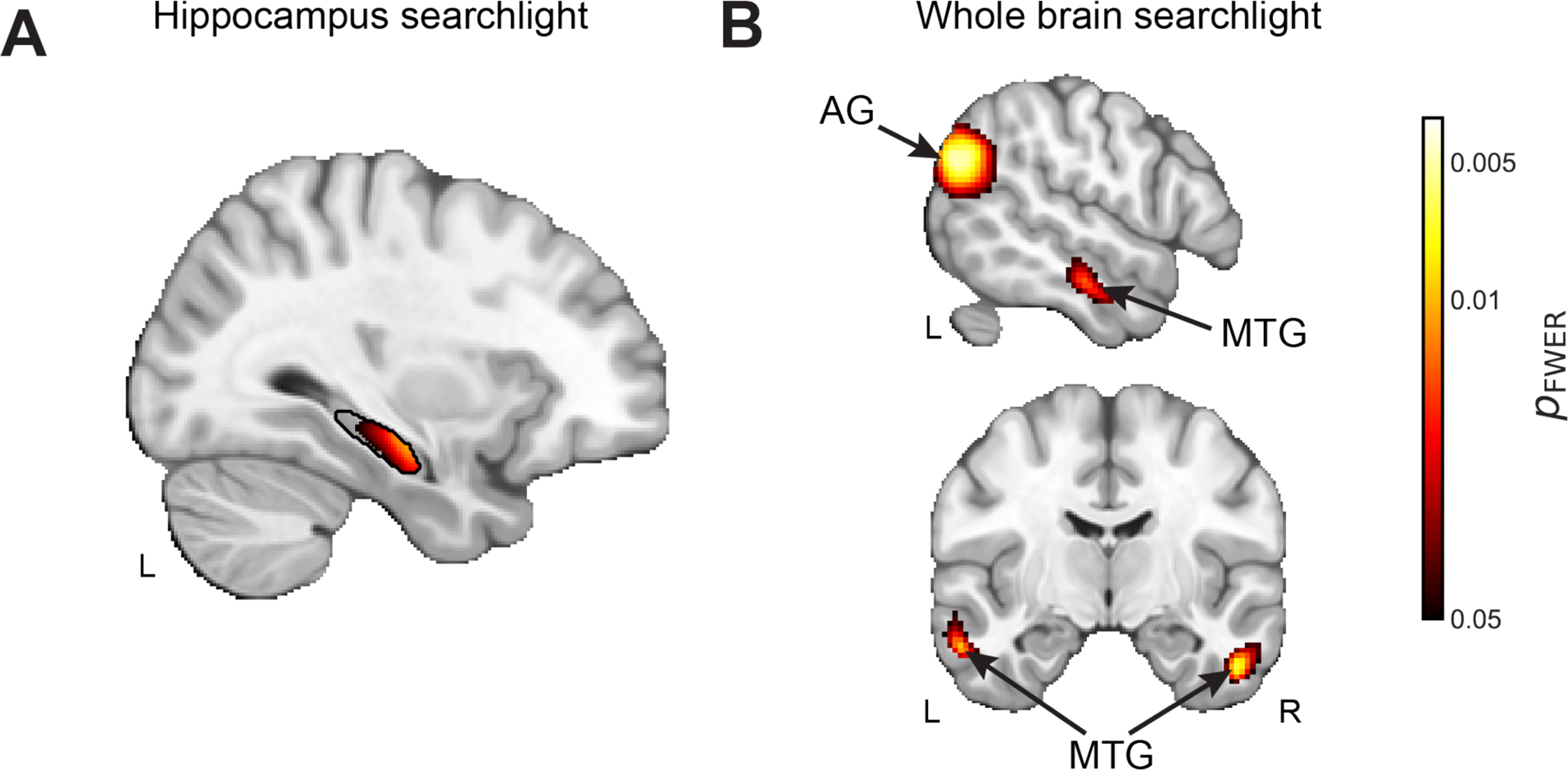
The left hippocampus represents affiliation and power decision trials differently. Decision trials of the same dimension had more similar neural patterns than trials of different dimensions. A) An analysis restricted to an anatomically defined hippocampus showed a significant cluster in the left hemisphere. B) An exploratory whole-brain analysis showed additional clusters in the bilateral middle temporal gyrus (MTG) and angular gyrus (AG). Participants were pooled across samples (n = 50); sample included as covariate. Permutation-derived p_FWER_ < 0.05.

After testing our hippocampus prediction, we ran a whole-brain exploratory analysis. This showed additional significant clusters, suggesting a distributed network represents these dimensions: the bilateral angular gyrus (right: peak voxel MNI x/y/z = 44/−57/35, cluster extent = 1824 voxels; left: peak voxel MNI x/y/z = −52/−66/29, cluster extent = 1303 voxels) and the bilateral middle temporal gyrus (right: peak voxel MNI x/y/z = 51/−7/−32, cluster extent = 327 voxels; left: peak voxel MNI x/y/z = −54/−7/−26, cluster extent = 574 voxels) Both the angular gyrus, a region situated within the temporo-parietal junction, and the middle temporal gyrus have been implicated in social cognition (e.g., theory of mind)^14,15^.

### The hippocampus represents affiliation and power locations within relationships

We have shown that the left hippocampus represents the affiliation and power trials differently, consistent with an abstract dimensional representation. Does it also represent the changing social coordinates of each character? To test this, we multiple-regression RSA searchlight to test whether left hippocampus patterns represent the characters’ changing social locations across interactions (see **Figure 3**). We restricted the distances to those from trial pairs from the same character and standardized the distances within character (see **Figure 3B-D**). We controlled for temporal distance to ensure the effect was not explainable by the time between trials, and for whether the trials shared the same underlying dimension (affiliation or power; see **Location similarity searchlight analyses** for more details). At the group level, we controlled for sample and the average reaction time difference between affiliation and power trials. Using the same testing logic as the dimensionality similarity analysis, we first tested our hypothesis in the bilateral hippocampus and found widespread effects in both the left (peak voxel MNI x/y/z = −35/−22/−15, cluster extent = 1470 voxels) and right (peak voxel MNI x/y/z = 37/−19/−14, cluster extent = 1953 voxels) hemispheres. The whole-brain searchlight analysis revealed additional clusters in the left putamen (−27/−3/14, cluster extent = 131 voxels) and left posterior cingulate cortex (−10/−28/41, cluster extent = 304 voxels).

**Figure 3.**
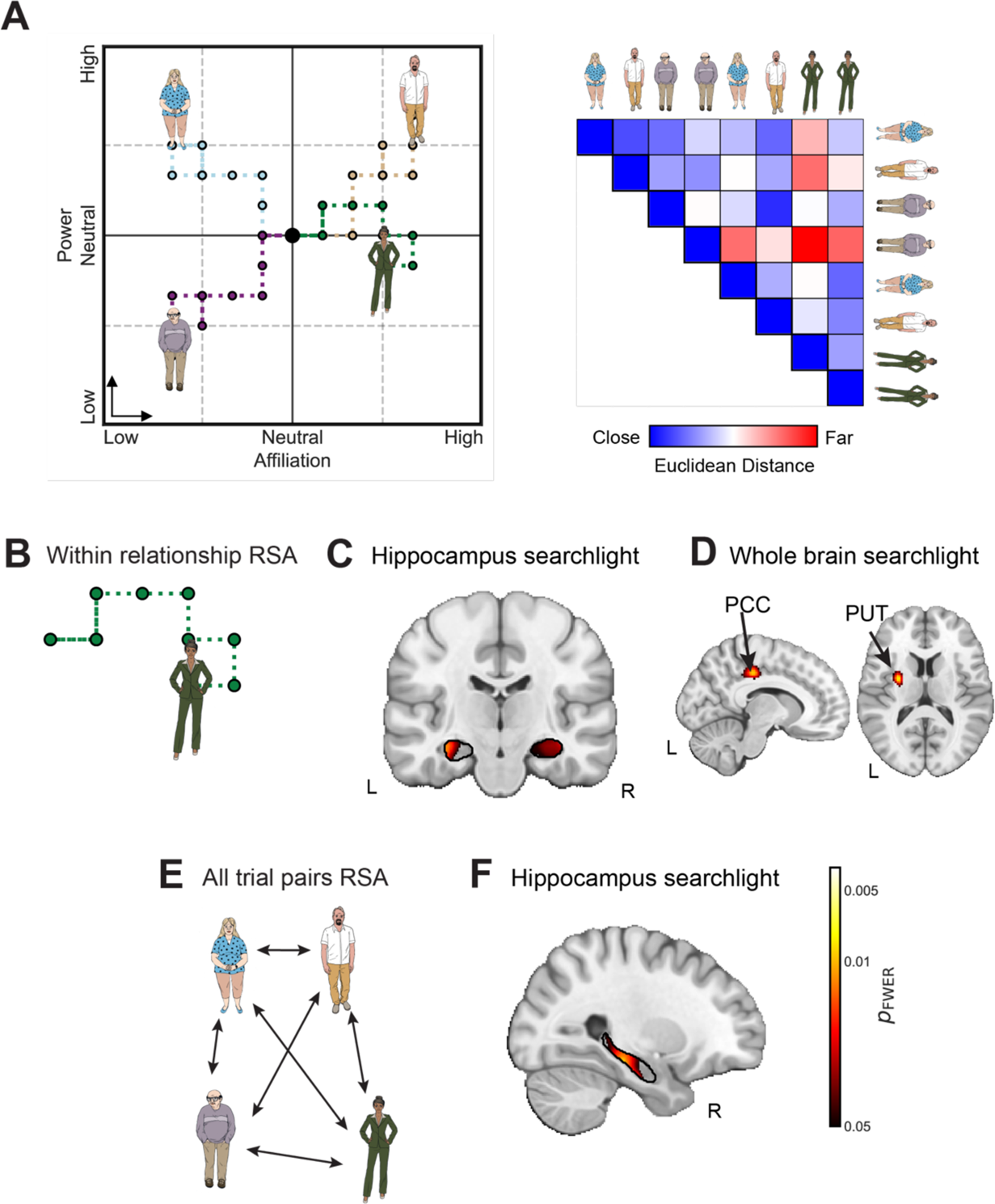
The hippocampus represents dynamically changing abstract social locations. A) We used representational similarity analyses (RSA) to study location-like effects. We used the behavioral locations implied by the participants choices to compute behavioral distance matrices. The distance matrix shown is a small, toy schematic to represent the logic; there were 60 decision trials in the task. We ran regressions and included control distances for task- and time-related variables to isolate the contribution of abstract location-related distances. B) We ran a within-character version of the RSA, where distances were within-character, to test for the brain dynamically tracking locations over relationship-specific social interactions. C) The anatomically defined left and right hippocampi both showed within-character location-like effects, in a region-of-interest analysis. The left show effects from the mid- to posterior hippocampus, whereas the right was mainly in mid-hippocampus. D) An exploratory whole-brain analysis also showed clusters in the posterior cingulate cortex (PCC) and putamen (PUT), as well as a small cluster in the right inferior temporal cortex (not shown). E) We ran an additional RSA, including all trial pairs, to test for evidence of a shared map-like representation. F) In a hippocampal specific analysis, there was an effect in the mid-to-posterior right hippocampus. In these analyses, participants were pooled across samples (n = 50); sample included as covariate. Permutation-derived p_FWER_ < 0.05.

We then asked a second, complementary question: does the hippocampus represent all interactions, across characters, within a shared map? To test for this map-like structure, we repeated the analysis but now included all trial pairs, z-scoring distances globally rather than within character (**Figure 3E-F**). The remainder of the procedure followed the same logic as the preceding analysis. The hippocampus analysis revealed an extensive right hippocampal cluster (27/−27/−14, cluster extent = 1667 voxels). The whole-brain analysis did not show any significant clusters.

## Discussion

Here, we study a case of conceptual trajectories, which are sequences of abstractly, rather than physically, related events, and whose order is meaningful rather than arbitrary or physically determined (spatial or temporal sequences). We show that by connecting conceptually related interactions, the hippocampus represents evolving social relationships as sequences of neural states.

We first show that left hippocampal patterns are more similar in social interactions with the same underlying dimension (i.e., affiliation compared to affiliation, and power compared to power) than interactions with different dimensions (i.e., affiliation to power), consistent with abstract affiliation and power representations. Then, we show that the hippocampus tracks the changing social locations (affiliation and power coordinates), above and beyond the effects of dimension or time; the hippocampus seemed to reflect both the changing within-character locations, tracking their locations over time, and locations across characters, as if in a shared map. Thus, these results suggest that the hippocampus does not just encode static character-related representations but rather tracks relationship changes in terms of underlying affiliation and power.

Previous work has shown that the hippocampus tracks temporally sequential events, such as trajectories in physical space (i.e., traversing sequentially connected locations^16^) as well as sequences of arbitrary information units, such as ordered lists of words or scenes (e.g.,^17^). Social relationships, however, are *conceptual* sequences: the events that constitute a relationship are spatially and temporally distant but conceptually linked, constituting a trajectory in abstract social space. Stacking the neural representations of these interactions together, then, would constitute the neural trajectory of the relationship through social space.

Why would the hippocampus encode abstract sequences with a neural trajectory? We cannot answer this question directly with fMRI, but there is a plausible account from the neurophysiological literature. Hippocampal neurons are known to track the physical locations of oneself^18^ and others^19^, as well as social identity^20^ and non-spatial sensory information^21^, suggesting the hippocampus may compute locations in abstract spaces—including social space. Moreover, the activity of hippocampal cell populations is ordered with respect to task dimensions, such that population-level patterns track locations in task space^21,22^. Because hippocampal firing fields overlap to cover the task space^23^, trajectories through task space should activate sequences of correlated hippocampal patterns, forming a neural trajectory that reflects the behavioral trajectory. This activity may be intrinsically low-dimensional: many hippocampal cells co-activate to encode locations in task space, restricting the possible activity patterns to a low-dimensional subspace^24^. Many possible neural states in theory become many fewer in practice, as neuronal correlations restrict intrinsic activity to low-dimensional manifolds. In the case of conceptually related events that are separated in time, space, and context—as in the case of episodes of social interactions over a lifetime—it is possible that at each interaction, the hippocampus retrieves a representation of the last interaction’s social location and then infers how the current interaction changes the relationship’s location.

The approach used in this study can be applied to similar questions in non-social domains. For example, context-related effects^25^ may also be explained in a similar manner: memories of different episodes that share a higher-order context could have correlated patterns of activity, creating trajectories through a conceptual space. The structure of the manifold that these trajectories evolve along may hint at its underlying dynamics, allowing the formation and testing of novel computational hypotheses.

In summary, we provide evidence that the hippocampus represents social relationships with ordered sequences of neural patterns. Each relationship had its own sequence of states, and all relationships were embedded in the same manifold. These results put forward a novel way to look at representations of all kinds of evolving latent relationships, physical to social.

## Methods

### Independent samples

To ensure the effects we find are robust, we tested our hypotheses in two independent samples, called the “Initial” and “Validation” samples. The Initial sample was collected in a previous study^3^ and included 18 of 21 participants (8 female), after excluding 1 for psychiatric evaluation (Psychiatric Diagnostic Screening Questionnaire) and 2 for DICOM corruption during archiving. The Validation sample included 32 of 39 participants (17 female), after excluding 2 for poor post-task character memory (<= chance [20%] for 5 main characters) and 5 for high motion (mean framewise displacement >= 0.3mm). The two samples did not differ in terms of participant sex (χ^2^ = 0.45, *p* = 0.5) but were different in age (in years; Initial: mean (*M*) = 29.33, standard deviation (*SD*) = 3.42; Validation: *M* = 44.42, *SD* = 9.69; Welch’s unequal variances t-test *p* < 0.001).

The Institutional Review Board of the Icahn School of Medicine at Mount Sinai approved the experimental protocols for both samples. All participants provided written informed consent and were paid for their participation.

### Social Navigation Task

#### Task description

The Social Navigation Task is a narrative-based social interaction game. Pre-task instructions are minimal: participants are told they will complete a task where they interact with different fictional characters, and to just behave as they naturally would. Importantly, the participants are never told or otherwise taught about the affiliation and power dimensions underlying the interactions. From the point-of-view of the participant, they are simply interacting with characters in a narrative and the affiliation and power dimensions are fully latent (i.e., unobserved or implicit).

There were 6 characters; their gender and racial presentations were counterbalanced. Gender was flipped across two versions (three men and three women), and in the Validation sample, the characters’ skin color was also counterbalanced with lighter- and darker-skinned characters variants of the same base images. The task version used for the Initial sample had approximately the same text and options as the Validation sample’s version (with small modifications), but with more cartoon-like character images^6^.

The narrative begins with the participant being told that they have just moved to a new town, and they need to find a job and a place to live. It unfolds over various social interactions that vary in their details and then culminates in the participant having successfully found employment and housing. There are two types of trials: “narrative” trials where background information is provided or characters talk or take actions (a total of 154 trials), and “decision” trials where the participant makes decisions in one-on-one interactions with a character that can change the relationship with that character (a total of 63 trials). On each decision, participants used a button response box to select between the two options. The options (1 or 2, assigned to the index and middle fingers) choice directions (+/−1 arbitrary unit on the current dimension) were counterbalanced.

The sequence of trials, including both narrative and decision trials, were fixed across participants; all that differs are the choices that the participants make. Narrative trials varied in duration, depending on the content (range 2-10 seconds), but were identical across participants. Decision trials always lasted 12 seconds, with two options presented until the participant made a choice, after which a blank screen was presented for the remainder of the duration. All six characters’ decision trials are interleaved with one another, and with the narrative slides. On average, after a decision trial for a given character, participants view ~11 narrative slides and complete ~3 decisions for other characters before returning to another decision with the same character, such that each character’s choices are separated by an average of ~20 seconds (ranging from 12 seconds to 10 min).

#### Affiliation and power decisions

Decision trials were categorized as either affiliation or power decisions. Each decision trial presented two options, one increasing (+1) and one decreasing (−1) the character’s position along the relevant dimension. Five main characters each had 6 affiliation and 6 power decisions, and a sixth neutral character had 3 neutral decisions (i.e., where the choices were inconsequential; not used for analysis). Affiliation decisions were defined as decisions whether to share physical touch, physical space or information (e.g., to share their thoughts on a topic). Power decisions were defined as decisions to issue or submit to a directive/command, or otherwise exert or give control. Examples of both trial types are shown in **Table 1**.

**Table 1.** Examples of affiliation and power interaction decisions. Several examples of affiliation and power interactions are shown. The slide preceding the choice trial is shown to give the context for the interaction, along with the decision that would increase and the decision that would decrease the location along the relevant dimension.

| Decision dimension | Previous slide context | Increase option | Decrease option |
| --- | --- | --- | --- |
| Affiliation | Chris goes in for a hug. | You hug him for a long moment. | You shake his hand instead. |
| Affiliation | Anthony is taking the elevator downstairs with you. | You: "Looks like you're the real person running this place - everybody really relies on you!" | You check your email on your phone: "Is it always that hectic in here?" |
| Affiliation | Mrs. Newcomb gets a phone call. Something is wrong; she seems very worried. | You feel concerned: "Everything alright, Jane? What is it?" | You don't want to get involved: "I'll give you privacy. I think we're done, anyway." |
| Power | Maya: "It doesn't matter if you're late. We're grabbing coffee." | You: "I can't - gotta go. Maybe tomorrow." | You really don't want to be late but reluctantly agree to grab coffee. |
| Power | Kayce is approaching the car. Chris gestures for you to move over to the middle seat. | You wait for Chris to take the middle seat. | You immediately move over to the middle seat. |
| Power | Anthony: "By the way, I have a lead on a place. It's a condo with two separate units." | You: "Get me the contract and floor plan and tell them you need until tomorrow." | You: "Let me know when you have more information." |

To ensure that any observed neural or behavioral differences were not confounded by trivial features of the text, we tested for differences between the affiliation and power trials (where the two options are concatenated). There were no differences in word count (affiliation average = 26.6, power average = 25.6; t-test *p* = 0.56). The text’s sentiment also did not differ between these trial types (t-test *p* = 0.72), as quantified by comparing sentiment compound scores (from most negative, −1, to most positive, +1), using a Large Language Model (LLM) specialized for sentiment analysis^26^.

Our framework assumes that affiliation and power trials differ in their semantic content–that is, in the conceptual meaning of the text, beyond word count or sentiment. To test this assumption, we used an LLM-based semantic embedding analysis. Each decision trial was embedded into a semantic vector. We then measured the cosine similarity between pairs of trials and calculated the difference between average within-dimension similarity (affiliation-affiliation and power-power comparisons) and average between-dimension similarity (affiliation-power comparisons) and assessed its statistical significance with permutation testing (1,000 shuffles of trial labels). As expected, decision trials of the same dimension were more similar to each other than trials of different dimension, across multiple LLMs (OpenAI’s text-embedding-3-small^27^: similarity difference = 0.041, *p* < 0.001; all-MiniLM-L12-v2^28^: similarity difference = 0.032, *p* < 0.001).

### Post-task measures

#### Task memory

To validate that the participants attended to the task, participants answered 30 memory questions about the characters after the task was completed (outside the scanner). In each question, the options were 5 of the 6 characters (including the neutral): each character was presented as an option in 25 questions and was the correct answer in 5 of those questions.

#### Self-reported character placement

Participants may form a subjective representation of the characters’ 2D social locations. To probe this, the participants completed a character placement task outside of the scanner after the task (Validation sample only). The participants were instructed to drag-and-drop colored dots representing each of the 6 characters onto locations in a 2D affiliation and power space, according to the participant’s perceptions of their relationships with the characters. The placements were relative to the participant’s theoretical point-of-view, which was represented by a red dot placed at the max affiliation and neutral/middle power values.

### Behavioral analysis

#### Behavioral modeling

Each character started the task at the neutral origin (0, 0) of the social space. With each participant choice, the current character’s coordinates were implicitly updated in the positive or negative direction along the current interaction dimension (i.e., each decision moved the character +/− 1 arbitrary unit along affiliation or power). For any given decision trial (t), the current character’s (c) affiliation and power coordinates are the cumulative sums of the trial-wise affiliation and power decisions:

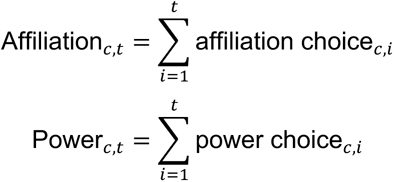

where choice value is −1 or +1 if the trial is an interaction of that dimension, and 0 if not. Thus, each participant’s sequence of interaction choices create a relationship trajectory through the latent affiliation and power space (see **Figure 1**).

#### Self-reported character placement analysis

We modeled the choices in the character-specific interaction sequences as affiliation and power coordinates; if the brain represents these coordinates, participants may have subjective access to them. Thus, we expected participants’ post-task placements of the characters into a 2D affiliation and power space to be closer to their end-of-task behavioral locations than expected by chance. For each participant in the Validation sample, we calculated the mapping error as the average Euclidean distance between the character-wise behavioral and self-reported locations. To establish participant-specific chance distributions, we permuted the subjective locations and re-calculated this error 100 times for each participant. We compared each participant’s average mapping error against their chance distribution by calculating a z-score with respect to the permutation distribution. We then tested whether these mapping errors were smaller than 0 with a left-tailed t-test.

Given that retrieving the subjective locations should be memory-dependent, we also predicted a negative relationship between mapping error and post-task memory recall: participants with better memory of the interactions should place the characters closer to their behavioral locations. We tested this prediction using Pearson’s correlation and a left-tailed p-value.

### fMRI acquisition and pre-statistics

#### Image acquisition

Scans were collected in a single run of approximately 26 minutes. Both samples’ data were acquired on 3-Tesla (3T) scanners but with different image acquisition parameters.

##### Initial sample

Images were collected on a Siemens Allegra 3T scanner (Siemens, Erlangen, Germany). T2*-weighted images were collected with a single-shot echo-planar imaging (EPI) pulse sequence with the following parameters: flip angle = 90°, echo time (TE) = 35 milliseconds (ms), repetition time (TR) = 2 seconds (s), 36 slices, 64 x 64 matrix, voxel size = 3 millimeter^3^ (mm^3^). T1-weighted images were collected with a magnetization-prepared rapid gradient-echo (MPRAGE) protocol with voxel size = 1 mm^3^.

##### Validation sample

Images were collected on a Siemens Skyra 3T scanner. T2*-weighted images were collected with a multiband slice EPI pulse sequence with the following parameters: multiband acceleration factor = 7, flip angle = 60°, TE = 35 ms, TR = 1 s, 70 slices, 108 x 108 matrix, voxel size = 2.1 mm^3^. T1-weighted images were collected with a MPRAGE protocol with voxel size = 0.8 mm^3^.

#### Image preprocessing

Image preprocessing was conducted using SPM12 (Wellcome Trust Centre for Neuroimaging), with standard steps. To correct for head motion, the functional images were realigned to the first volume using 6 parameter rigid body transformation (3 translations and 3 rotations) and then unwarped to account for magnetic field inhomogeneities. The realigned images were slice-time corrected to the middle slice using normalized mutual information, then co-registered in alignment with the MPRAGE to the mean unwarped image. The MPRAGE image was then segmented into 6 tissue classes (gray matter, white matter, cerebral spinal fluid, skull, soft-tissue and air) and the resulting forward deformation parameters were used to normalize the images to a standard Montreal Neurological Institute (MNI) template image (using 4th-degree B-spline interpolation). The functional images were not smoothed.

### fMRI analysis

#### Trial-wise general linear modeling

To estimate single trial activation patterns, general linear models (GLMs) were fitted to each voxel in the functional images using SPM12. Unsmoothed images were used to preserve the spatial resolution of the multi-voxel patterns. Microtime resolution was set to the number of slices collected and microtime onset was set to the middle slice. To reduce temporal autocorrelation in the time series, we used a high-pass filter of 128 seconds (1/128 Hertz) and prewhitening (with SPM’s FAST algorithm).

The design matrices had separate regressors for each decision trial to estimate all decision trials’ weights in the same regression (i.e., a least-squares-all approach). For different analyses (see below), we modeled the events with a stick function at trial onset or boxcar function over the reaction time (from onset to the button press), depending on whether the neural representation was assumed to be independent of the decision. The narrative trial onsets were modeled by an additional regressor. The decision and narrative trial regressors were convolved with a canonical hemodynamic response function to model the blood oxygenation level dependent (BOLD) signal changes expected from the events. The six realignment parameters from preprocessing were included in the design matrices to regress out residual motion-related variance. To further improve the signal-to-noise ratio, the GLMs were only estimated for voxels with an average signal greater than 50% of the global signal. After GLM estimation, each participant had a beta image with a beta series (β_1_, β_2_, …, β_63_) for each voxel that reflects trial-specific activation magnitudes. The Initial sample’s beta images were resampled to match the higher spatial resolution Validation sample’s images, so that all analyses cover the same number of voxels and amount of brain volume across participants.

#### Social dimension similarity searchlight analysis

The affiliation and power dimensions are akin to social contexts: we expected the hippocampus to represent interactions of the same dimension more similarly than interactions of different dimensions. In other words, we expected the affiliation and power interactions to be represented abstractly, such that their representations generalize across trials. We used a representational similarity analysis (RSA) searchlight to test this hypothesis (see **Searchlight analysis details** below). We assumed that representations of social dimension are independent of the decisions themselves, and so we used participant-specific trial-wise beta images where each decision trial was modeled with a stick function regressor at the trial onset. We expected the same dimension trial pairs to have more correlated neural patterns than different dimension trial pairs. For each participant and each searchlight sphere, we calculated the difference between the average neural pattern correlations (Pearson’s *r* with Fisher’s z-transform) for same dimension trial pairs versus different dimension trial pairs; the resulting statistic was stored in the center voxel.

Importantly, this analysis naturally balances various narrative-related variables. As described in the methods (see **Affiliation and power decisions**), affiliation and power trials were not different in word count or sentiment. Character identity is also balanced, because the characters appear in the same number of affiliation and power trials. Further, the number of previous decision trials with specific characters were not statistically different between same and different dimension trial pairs (t-test *p* = 0.55). However, there was a significant difference in the average reaction time between affiliation and power decisions across participants (t_49_ = 6.92, *p* < 0.001; affiliation mean = 4.92 seconds (s), power mean = 4.51 s), so we controlled for this in the group-level analysis.

#### Location similarity searchlight analyses

We also expected the hippocampus to represent the different characters’ changing social locations, which are implicit in the participant’s choices. We used multiple regression searchlight RSA to test whether hippocampal pattern dissimilarity increases with social location distance, based on participant-specific trial-wise beta images where boxcar regressors spanned each trial’s reaction time.

We ran two complementary regression analyses to address two related questions. First, we asked whether the hippocampus represents how a specific relationship changes over time. For this analysis, for each participant and each searchlight, we computed character-specific (i.e., only for same character trial pairs) correlation distances between trial-wise beta patterns and Euclidean distances between the social location behavioral coordinates. Distances were z-scored within character trial pairs to isolate character-specific changes. The second analysis asked whether the there is a common map-like representation, where all trials, regardless of relationship, are represented in a shared coordinate system. Here, we included all trial pairs and z-scored the distances globally.

For both regression analyses, we included control distances to control for possible confounds. To account for generic time-related changes, we controlled for absolute scan-time difference, as this correlated with location distance across participants (see **Temporal autocorrelation of hippocampal beta patterns** in the supplement). Although the square of this temporal distance did not explain any additional variance in behavioral distances, we ran a robustness analysis including both temporal distance and its square and saw qualitatively the same clusters with similar effect sizes. As such, we report the main analysis only. We included binary dimension difference (0 = trial pairs of different dimension, 1 = trials pairs of the same dimension), to ensure effects could not be explained by dimension-related effects. In the group-level model, we controlled for sample and the average reaction time between affiliation and power decisions.

#### Searchlight analysis details

Each searchlight was a sphere with a diameter of 11 voxels. We smoothed the participant-level searchlight images with an 8mm Full Width at Half Maximum (FWHM) kernel and used a group-level voxel-wise group-level General Linear Model, controlling for sample and the average reaction time difference between affiliation and power decisions. We used right-tail tests to test whether the hypothesized effect had a positive relationship to neural pattern similarity. The family-wise error rate (FWER) for significant testing was controlled at an alpha of 0.05, using a max-type permutation testing procedure^29^.

#### Region-of-interest details

In our searchlight analyses, we specifically tested the bilateral hippocampus^6,10^. We defined left and right hippocampal masks from the Harvard-Oxford Atlas, co-registered and re-sampled them to the Validation sample’s functional images and then binarized at a 50% probability threshold. We used small volume correction in the hippocampi for our main tests. For whole-brain exploratory analyses, we corrected for the total number of (non-hippocampal) comparisons, and thresholded so that clusters were at least 50 contiguous voxels.

## Acknowledgments

Funding: DS is supported by the National Institute of Health, USA (R01MH122611, R01MH123069); KB is supported by the National Institute of Drug Abuse, USA (K23-DA045928); MS is supported by the National Institute of Mental Health, USA (F31MH123123).

## Author contributions

MS and DS conceived of the study. PKB, VS and KB collected the data. MS ran the analyses. MS, DS and KB wrote the manuscript.

