## Supplemental file for "Neural Trajectories of Conceptually Related Events"

### Supplement

#### Temporal autocorrelation of hippocampal activity

The Social Navigation Task is a narrative-based task, where the relationships with characters evolve over time; trial pairs that are close in time may have more similar fMRI patterns for reasons unrelated to social mapping (e.g., slow drift). It is important to account for the role of time in our analyses, to ensure effects go beyond simple temporal confounds, like the time between decision trials. To aid in this, we quantified how fMRI signals change over time using a pattern autocorrelation function across decision trial lags. We defined the left and right hippocampus and the left and right intracalcarine cortex using the Harvard-Oxford atlas and thresholded them at 50% probability. We chose intracalcarine corex as an early visual control region that largely corresonds to primary visual cortex (V1), as it is likely to be driven by the visually presented narrative. We used the same trial-wise beta images as in the location similarity RSA (boxcar regressors spanning each decision trial's reaction time). For each participant and region-of-interest (ROI), we extracted the decision trial-by-voxel beta matrix and quantified three kinds of temporal dependence: beta autocorrelation, multivoxel pattern correlation and multivoxel pattern correlation after regressing out temporal distance.

To estimate the temporal autocorrelation of the trial-wise beta values, we treated each voxel's beta values as a time series across trials and measured how much a voxel's response on one trial correlated (Pearson) with its response on previous trials. We averaged these voxel wise autocorrelations within each ROI. At one trial apart (lag 1), both the hippocampus and V1 showed small positive autocorrelations, indicating modest trial-to-trial carryover in response amplitude (see **Supplemental figure 1**) that by three trials apart was approximately 0.

Because our representational similarity analyses depend on trial-by-trial pattern similarity, we also estimated how multivoxel patterns were autocorrelated over time. For each lag, we computed the Pearson correlation between each trial's voxelwise pattern and the pattern from the trial that many trials earlier, then averaged those correlations to obtain a single autocorrelation value for that lag. At one trial apart, both regions showed positive autocorrelation, with V1 having greater autocorrelation than the hippocampus; pattern correlations between trials 3 or 4 trials apart reduced across participants, settling into low but positive values. Then, for each participant and ROI, we regressed out the effect of absolute trial onset differences from all pairwise pattern correlations, to mirror the effects of controlling for these temporal distances in regressions. After removing this temporal distance component, the short lag pattern autocorrelation dropped substantially in both regions. The similarity in autocorrelation profiles between the two regions suggests that significant similarity effects in the hippocampus are unlikely to be driven by generic temporal autocorrelation.

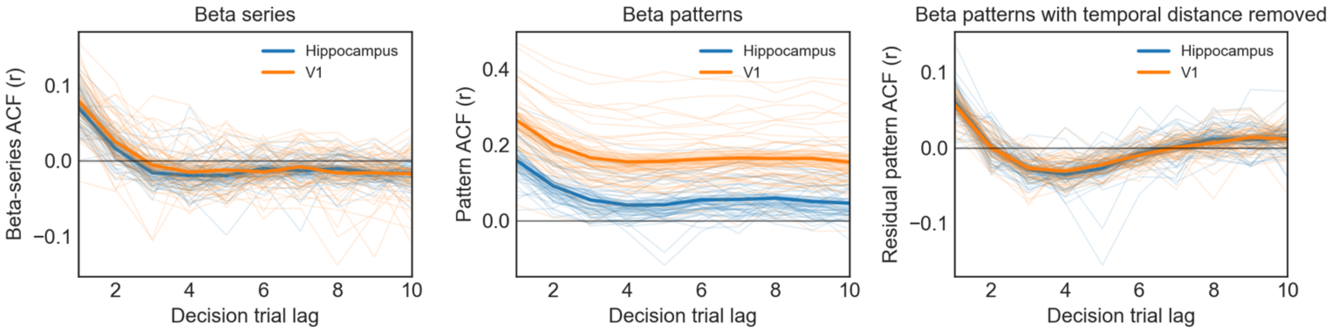


***Supplemental figure 1.*** *For left and right hippocampus and a region in primary visual cortex (V1), we plotted decision trial lag autocorrelation curves for (1) voxel-wise betas, (2) multivoxel beta patterns, and (3) multivoxel beta patterns after regressing out temporal distance. Because hemispheres showed very similar curves within participant, we averaged across left and right ROIs for visualization and show individual-participant curves, and the mean across participants for each lag.*

#### Relationship between behavioral location distance and temporal distance

We also quantified how temporal distances between trials relates to their behavioral location distances, participant by participant. Our dimension similarity analysis controls for temporal distance between trials by design (see **Social dimension similarity searchlight analysis**), but our location similarity analysis does not. To decide on covariates to include in the analysis, we tested whether temporal distances can explain behavioral location distances. For each participant, we computed the correlations between trial pairs' Euclidean distances in social locations and their linear temporal distances ("linear") and the temporal distances squared ("quadratic"), to test for nonlinear effects. We then summarized the correlations using one-sample t-tests. The linear relationship was statistically significant ($t_{49} = 12.24$, $p < 0.001$), whereas the quadratic relationship was not ($t_{49} = -0.55$, $p = 0.586$). Similarly, in participant specific regressions with both linear and quadratic temporal distances, the linear effect was significant ($t_{49} = 5.69$, $p < 0.001$) whereas the quadratic effect was not ($t_{49} = -0.20$, $p = 0.84$). Based on this, we included linear temporal distances as a covariate in our location similarity analyses (see **Location similarity searchlight analyses**), and verified that adding a quadratic temporal distance covariate does not alter the results. Thus, the reported location-related pattern similarity effects go beyond what can be explained by temporal distance alone.
